# Glycan Clock Forecasts Human Influenza A Virus Hemagglutinin Evolution

**DOI:** 10.1101/163345

**Authors:** Meghan O. Altman, Matthew Angel, Ivan Košík, Nídia S. Trovão, Seth J. Zost, James S. Gibbs, Scott E. Hensley, Martha I. Nelson, Jonathan W. Yewdell

## Abstract

Adaptive immunity to influenza A virus is limited by frequent mutations in the immunodominant head of hemagglutinin (HA). Over the last century, the upward trend in HA-head glycosylation indicates glycan addition can increase fitness, but its role in viral evolution remains unclear. Here, we report glycan evolution follows a clock-like rhythm, pacing the timeline, trajectory, and replacement of HA. Following pandemic introduction, glycans are added to HA at 4- to 6-year intervals, until a functional glycan limit is reached, after which, at 9- to 12-year intervals, glycans are either swapped between different sites, or the HA is replaced by a novel pandemic virus. Using this, we predicted the appearance of the newest glycan on pH1N1 HA. Phylogeographic reconstruction suggests these highly fit strains originated in the Middle East, before rapidly replacing all strains globally. Going forward, we can use this simple algorithm to forecast future glycan evolution and identify seasons with higher pandemic potential.

---

Seasonal influenza A virus (IAV) annually sickens millions and kills hundreds of thousands of people^1^. IAV remains endemic despite high levels of adaptive immunity in nearly all humans after early childhood. IAV persists due to the rapid selection of mutants with amino acid substitutions in the viral hemagglutinin (HA) glycoprotein that enable escape from neutralizing antibodies (Abs). HA is a homotrimeric glycoprotein with a globular head domain resting atop a stem that anchors HA to the virion surface. The head has a receptor binding site that attaches virions to cell surface terminal sialic acid residues, initiating infection. Ab binding to the head neutralizes virus by blocking attachment or the conformational alterations required for HA-mediated membrane fusion. Understanding Ab-driven HA evolution is essential to improving influenza vaccination, which currently, offers only moderate protection from infection^2,3^.

As newly synthesized HA enters the endoplasmic reticulum of IAV infected cells, oligosaccharides are attached to Asn residues present in the motif Asn-X-Ser/Thr-Y, where X/Y are all amino acids except Pro. Attached glycans can be high-mannose glycans or larger, more complex branched glycans^4,5^. HAs possess multiple highly conserved glycans on the stem domain crucial for HA folding and oligomerization^6^. Glycosylation of the head is much more variable, as evolving HAs add, retain, and occasionally lose potential head glycosylation sites^7,8^. Head glycans can promote viral fitness by shielding virus from antibody binding and tuning receptor affinity and specificity^9–12^. Conversely, glycans can deleteriously modulate receptor binding, enhance viral neutralization by innate immune lectins, or increase stress in the endoplasmic reticulum during translation^13–18^.

Glycosylation changes are generally not correlated with changes between antigenic clusters^19^, in part, because glycan addition typically lowers HA receptor avidity^20^, complicating binary assays such as hemagglutination inhibition (HI)^15^. Further, antigenic clusters are defined by HI assays performed using ferret anti-sera, but HA head glycans are selected to escape neutralization by human antibodies specific for antigenic sites that are weakly targeted by ferret antibodies^21^. Consequently, glycan evolution is typically unaccounted for in molecular evolution studies using HI-based analytic tools such as antigenic cartography^22,23^.

## Empirical analysis of relative HA size over time

Studies of HA glycan evolution rely nearly exclusively on bioinformatic predictions of glycan addition. NetNGlyc, the state of the art algorithm^24^, predicts N-linked glycosylation^5^, but provides neither a guarantee the glycans are added, nor information regarding the glycan structure. We therefore used gel electrophoresis to monitor glycan addition to HAs from 72 egg-grown H1N1 and H3N2 strains based on decreased HA migration (Fig. 1, Extended Data Fig. 1–4**, File 1, Table 1**). HA migration strongly correlates with the number of computationally predicted head glycosylation sites (Pearson coefficient r = 0.96, P < 0.01) (Fig. 1a). H1 adds an apparent 3.9 ± 0.1 kDa (R^2^ = 0.89) per glycan, suggesting the majority of such glycans are highly branched. The average H3 glycan, by contrast, is smaller (3.0 ± 0.2 kDa, R^2^ = 0.90), consistent with greater addition of simpler glycans. The size difference between H1 and H3 HAs is consistent with mass spectrometry characterization showing that H1 glycans skew to larger, more complex structures, while H3 glycans at several sites are majority high-mannose^5,9^.

**Figure 1.**
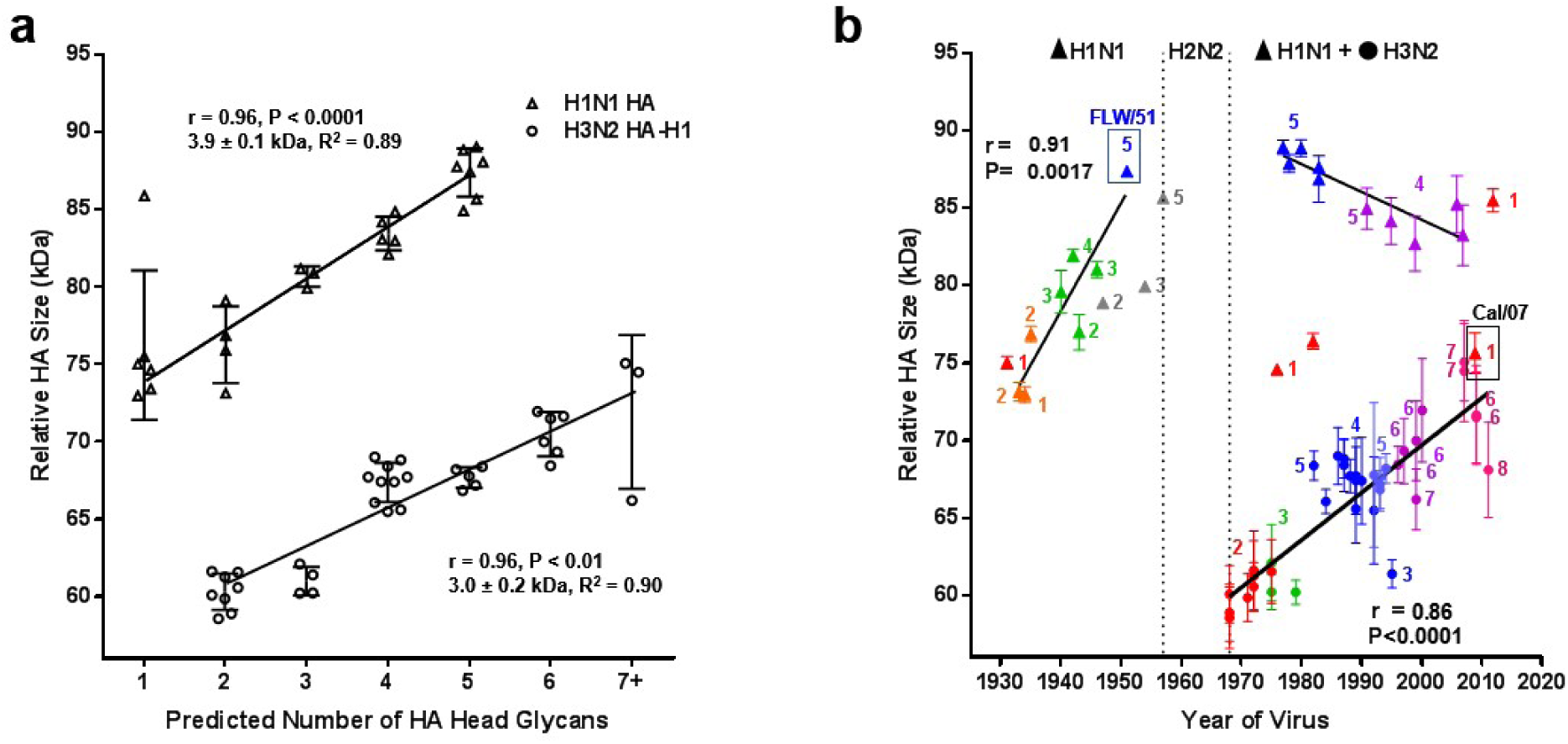
Quantitative survey of glycan evolution in IAV. **a,** Relative HA size, measured by migration rate through a gel, correlates with predicted number of head glycans. H1N1s (triangles) add 3.9 ± 0.1 kDa per glycan or 15.5 kDa total while increasing from 1 to 5 glycans. H3N2s (HA1, circles) add 3.0 ± 0.2 kDa per glycan or 15 kDa total while increasing from 2 to 7 glycans. **b,** HA size over time for H1N1s (triangles) and H3N2s (HA1, circles) reveals regular glycan addition. Strains are color-coded to match glycan groups in Extended Data Fig. 4. Numbers of HA-head glycans per strain are noted next to data. A glycan is added on average every 5 years for sH1N1 (3.9 kD) and every 8 years for H3N2 (3.0 kD). The strain most related to sH1N1 reintroduced in 1977, A/Fort Leonard Wood/1951, is boxed and labelled in blue. The pH1N1 strain, A/California/07/2009, is boxed and labelled in black. Pearson coefficient of correlation (r) and its P value are shown on both graphs. Error bars are standard error of the means from n = 3-5 blots.

Importantly, for H1 and H3 HAs, the total glycan mass added to the head during evolution in humans is, respectively, 15.5 kDa and 15 kDa. This is consistent with the idea of an upper limit of glycan shielding for H1 and H3 HA head domains totaling to ∼20 kDa (5 glycans for H1, 7 for H3). This is not strictly due to HA functional constraints: H1 HAs can accommodate at least 8 head glycans while maintaining *in vitro* fitness^25^. Rather it points to glycan-based fitness costs *in vivo*, possibly due to innate immune mechanisms or alterations in virus binding to receptors found in human airways^26^.

## H1 HA glycan evolution

Plotting relative HA size versus time reveals the history of seasonal H1N1 (sH1N1) glycosylation (Fig. 1b, triangles). First-wave sH1N1 (1933-1957) HAs behaved differently than the second-wave sH1N1 representatives (1977-2008). During the first-wave, relative HA size increased by an average of 3.9 kDa, or one glycan, every 5.6 years (slope = 0.69 kDa/year, R^2^ = 0.89).

As expected sH1N1 strains, that re-emerged after the 1977 re-introduction of sH1N1, exhibited similar relative HA size to their most common strain A/Fort Leonard Wood/1951 (inside blue box, Fig. 1b). Rather than increasing in number as occurred previously, electrophoresis confirms the NetNGlyc prediction that these 5 HA-head glycans strains were maintained until decreasing to 4 glycans after 1991.

These data indicate circulating H1 HAs did not acquire more than 5 head glycans. Indeed, HA sequences predicted to have 6 head glycans are highly unusual over the century of H1 evolution (1918-2017), except in 1986 and 1987, when they competed among H1 strains (Extended Data Fig. 5a). Rather than adding glycans, sH1N1 strains that reemerged after 1977 exhibited glycan adjustments spaced at 10.0 ± 0.9 year intervals (Fig. 2). In 1986, N71 pH1N1 (numbering convention is shown in **Extended Data Table 1)** replaced N172, as N144 swapped with N142, which though adjacent to 144, is spatially oriented to shield different epitopes. Around 1997, N286 was lost. Strains with the 4 remaining glycans circulated for 11 years before becoming extinct during the 2009 pH1N1 pandemic.

**Figure 2.**
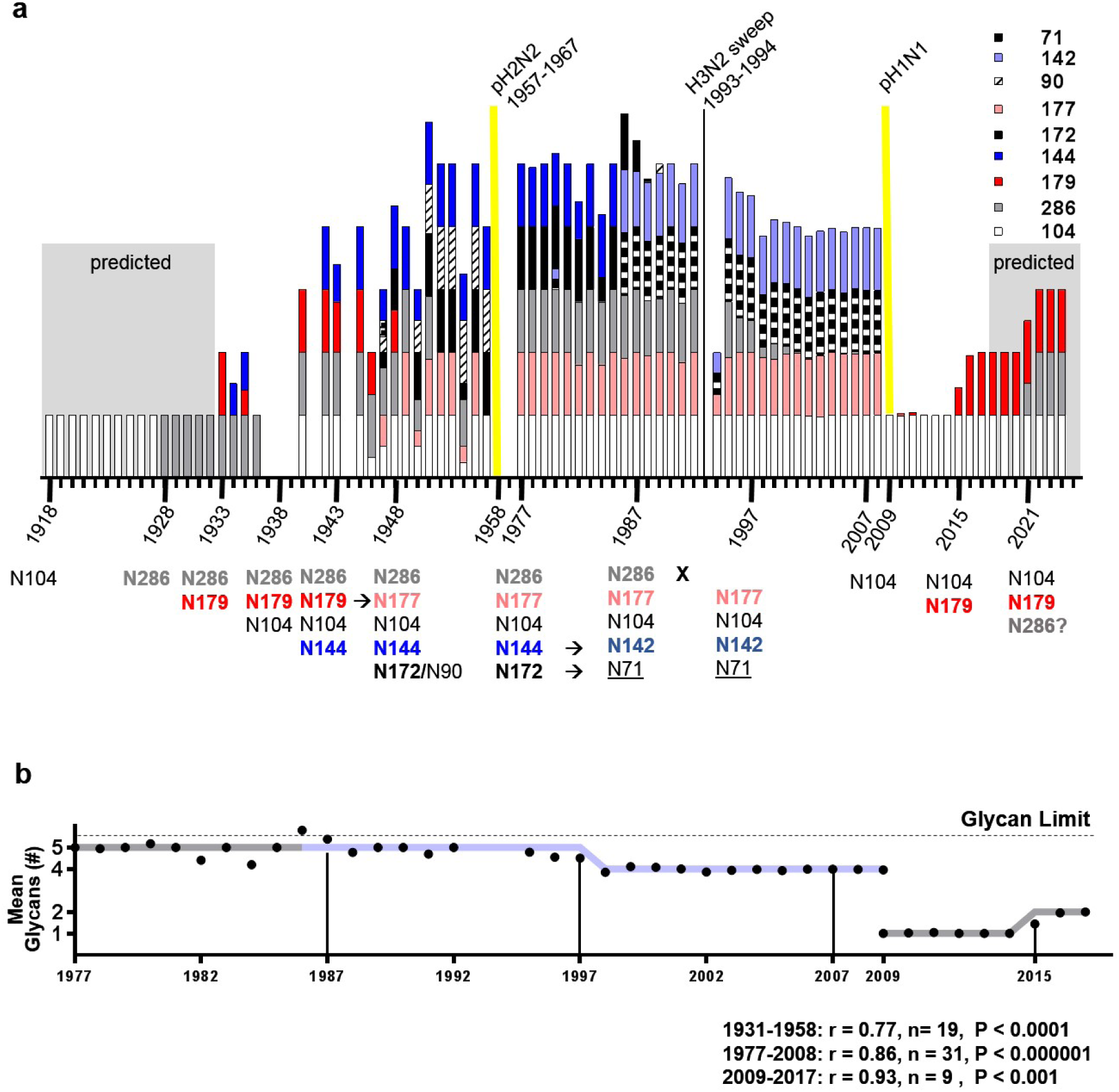
H1 Glycan Evolution 1918-2024. **a,** Stacked bars correspond to the percentage of all human H1N1 sequences from a given year containing a glycan listed in the legend. Dates of glycan transition are written on the timeline below along with the glycosylated residues. Projected values for strains from 1918-1933 and 2017-2024 are shown in shaded regions. Years with no data are left blank. Sequence counts for each year are shown in Extended Data Fig. 5. Pandemics are denoted by yellow lines. **b,** Mean predicted number of glycans among all sequences from each year are shown as black circles, which are displayed atop the model (grey line, light blue line indicates glycan swap from sequences containing N144/N172 to N142/N71). Black vertical lines denote predicted years for glycan evolution. Pearson’s coefficient of correlation (r) with sample number (n) and P value between the model and the data is shown below.

## H3 HA glycan evolution

As with H1 HAs, H3 HAs exhibit a regular rhythm of glycan evolution (Fig. 3). Original (1968) pandemic H3 strains had 2 head glycans. Four times since 1968, glycans were added to H3 HA at regularly spaced 5.0 ± 0.6 year intervals (N126 by 1974, N246/N122 by 1980, N133 + N122 by 1998, and N144 by 2004) (Fig. 4). This timing is similar to the average rate of glycan addition (one every 5.6 years) seen by SDS-PAGE for sH1N1 from 1933 to 1951.

**Figure 3.**
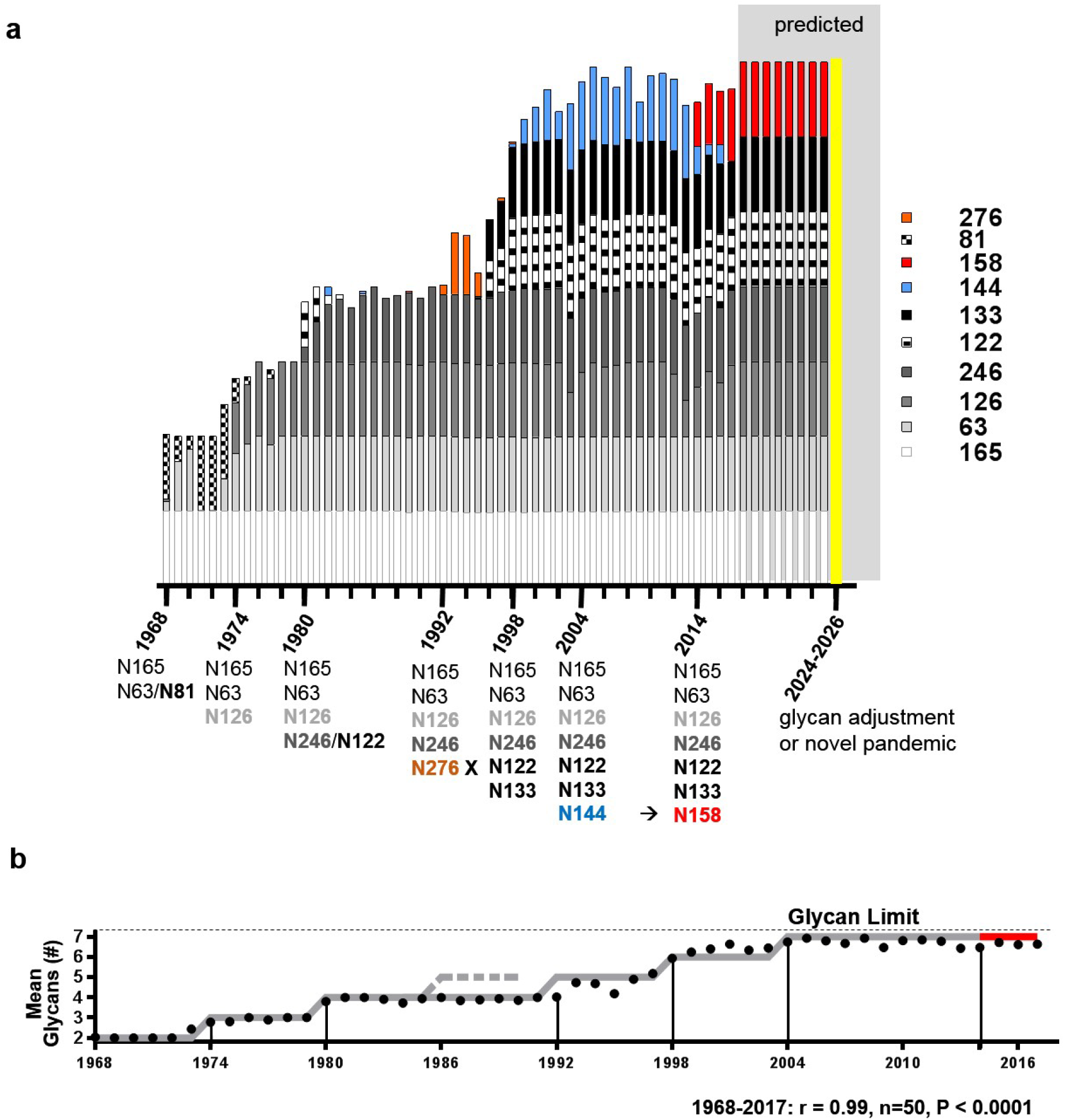
H3 Glycan Clock 1968-2026. **a,** Stacked bars correspond to the percentage of all human H3N2 sequences from a given year containing a glycan listed in the legend. Dates of glycan transition are written on the timeline below along with the glycosylated residues. Projected values for strains from 2017-2026 are shown in shaded regions. A potential pandemic is denoted by yellow line. **b,** Mean predicted number of glycans among all sequences from each year are shown as black circles, which are displayed atop the model (grey line, red line indicates glycan transition from sequences containing N144 to N158). Dashed grey line indicates when N276 would presumably have been added without functional constraint. Black vertical lines denote predicted years of glycan evolution from the model. Pearson’s coefficient of correlation (r) with sample number (n) and P value between the model and the data is shown below.

**Figure 4.**
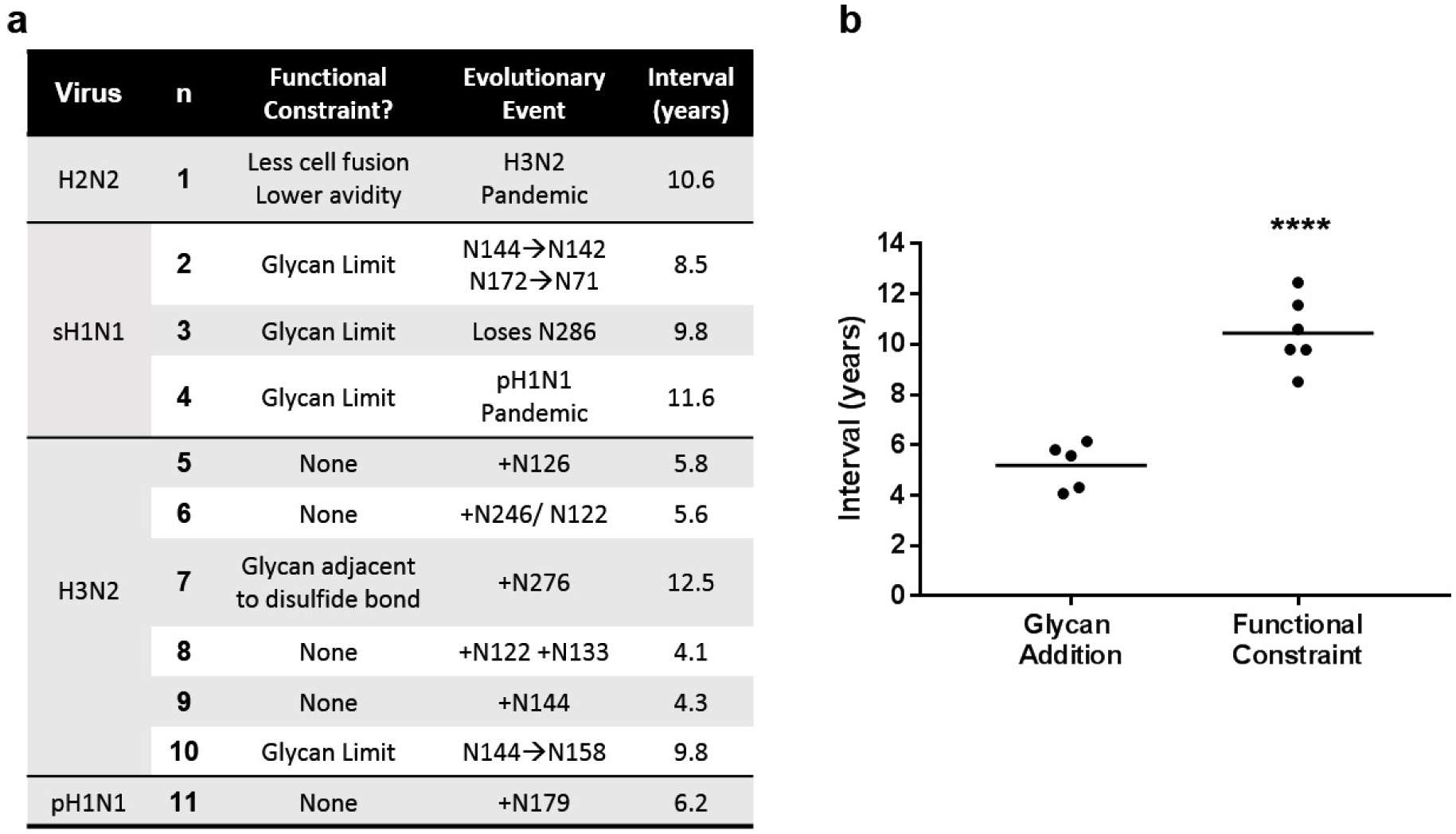
Quantitative analysis of evolutionary intervals. **a,** The interval (years) prior to occurrence of glycan altering events was calculated by fitting the proportion of changed sequences to a 3-parameter logistic regression and finding the inflection point of the curve. Events were classified as either having (n=6) or not having (n=5) a functional constraint that limited glycan addition. Starting dates were determined by a fixed sliding window every 10 years for sH1N1 and 6 years for H3N2 (black vertical lines in Figs. 2b and 3b). **b,** With no functional constraint, addition of a glycans occurs on average every 5.2 ± 0.9y. If the glycan is functionally constrained, glycan evolution occurs at 2-fold longer intervals, averaging 10.5 ± 0.9y (Student’s t-test ****P<0.0001).

With the addition of the 7th head glycan in 2004, H3 HAs had increased 15 kDa from the original 1968 strain (Fig 1b). It is apparent from the sequence data that H3 HAs with more than 7 head glycans lose in the competition for human circulation. While 8-glycan strains are present, they represent <1% of H3N2 sequences in any year (Extended Data Fig. 5b). Similar to second-wave H1 viruses, 9.8 years after reaching the glycan limit, a glycan swap occurred, as the glycan at residue N144 was replaced by a glycan at residue N158 (Extended Data Fig. 6a).

There was one exception in the 4- to 6- year interval pattern for H3 glycan addition. Addition of a fifth glycan at residue 276, occurred 12.5 years after the previous glycan addition (Fig. 3a, **orange**). In 1993 highly robust strains with N276 swept the globe, dominating sH1N1 strains, before disappearing after only 3 seasons. N276 is unique because the Asn residue bearing the glycan is immediately adjacent to a disulfide-bonded cysteine (C277). This site delineates the HA head from the HA stem. We speculate that the structural rigidity conferred by the disulfide reduced the fitness of N276, requiring epistatic changes elsewhere in the HA to improve competitiveness, both delaying fixation of the glycan addition, and also reducing its competitiveness against future strains lacking N276.

## H2N2 glycans did not evolve

Human H2N2 strains never added a glycan during their 11-year era (1957-1967). Notably, while selection of H2N2 strains with additional head glycans readily occurs *in vitro* under antibody pressure, these multi-glycan H2 HAs are functionally constrained, as they exhibit less cell fusion and lower receptor binding^27^. This is yet another example where functional constraints on glycan addition are associated with longer intervals between glycan change or pandemic replacement.

### Patterns for clock-like glycan evolution

Based on these observations we can identify 3 patterns that occur in human IAV evolution:

1. After introduction of a pandemic strain, glycans are typically added at intervals of 4- to 6-years.
2. Glycans are limited on the HA head to 5 and 7 glycans, respectively, for H1 and H3 HAs, with a total mass of ∼20 kDa due to preferred use of complex (H1) *vs.* simple (H3) glycans.
3. If glycan addition is constrained by fitness loss, then glycan adjustment (i.e. swapping positions) or pandemic replacement occurs at intervals of 9- to 12-years.

Quantitative analyses supporting the statistical significance of patterns one and three are shown in Fig. 4.

### Forecasting a glycan addition

Initial 2009 pH1N1 had a single predicted head glycan (N104). As expected, the HA of a representative early strain, A/California/07/2009 (Fig. 1b, inside black box), co-migrated with strains from the 1930s and two other swine-origin human strains, A/New Jersey/8/1976 and A/Memphis/4/1982 with single predicted head glycan (Fig. 1b, red triangles).

We initially deduced Pattern 1 solely from SDS-PAGE data (Fig. 1). From this, we predicted pH1, which appeared in 2009, should add a glycan ∼5.6 years after appearing. After identifying all potential HA head glycosylation sites in pH1 sequences, we found, as predicted, a second glycan site arose through an S to N change at residue 179 over the 2015-2016 season (Fig. 5a).

**Figure 5.**
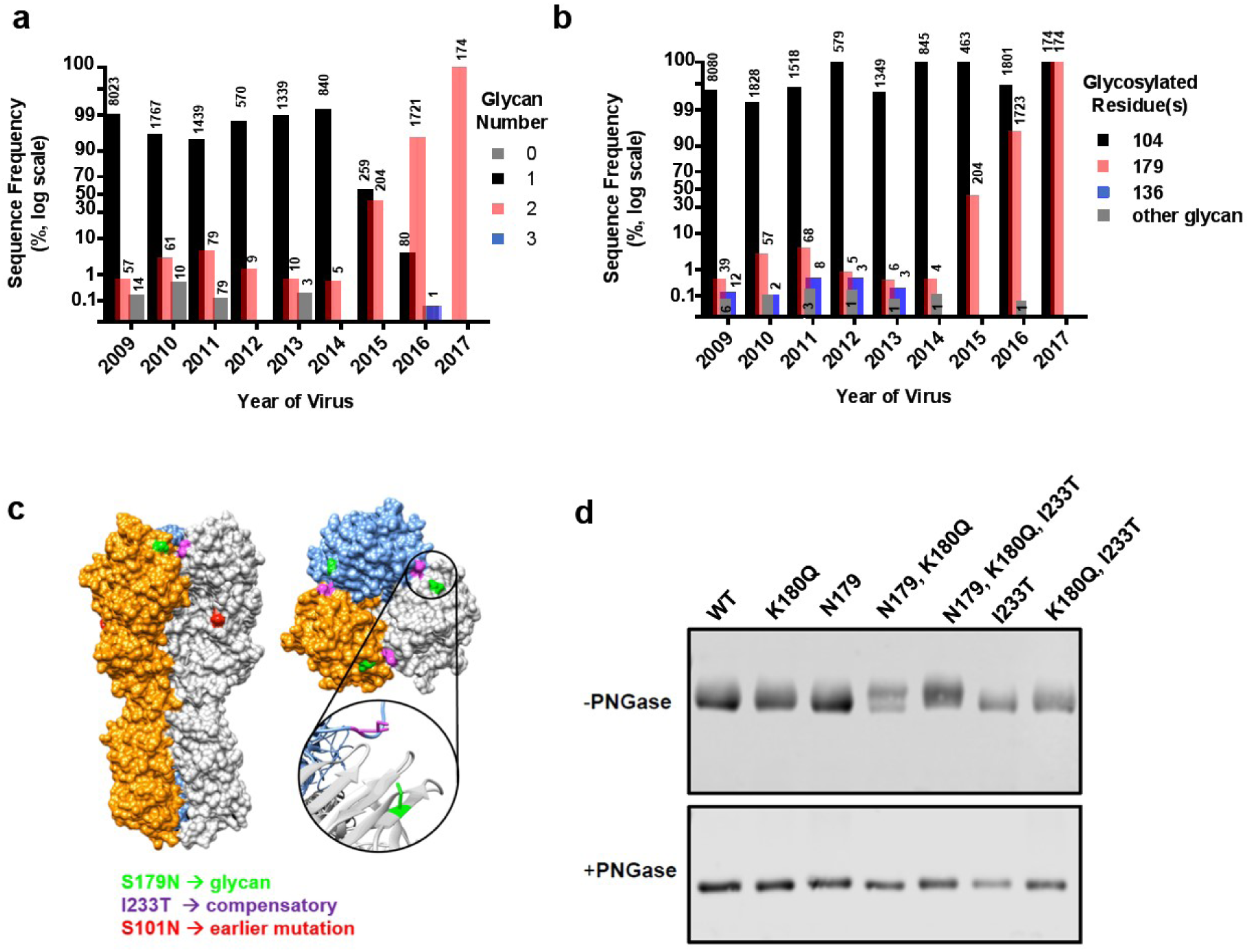
Forecasted pH1N1 glycan addition. **a,** Frequency of human pH1N1 sequences in FluDB by number of predicted glycans in each calendar year. Number of sequences represented by each bar is shown on top of bar. Until January 2015, >90% of sequences had 1 glycan (black). Starting in 2015, strains with 2 glycans swept (red). **b,** Frequency of individual residues glycosylated in human pH1N1 sequences. N104 predominates throughout (>95%, black). N179 (red) and N136 (blue) appear at low frequencies until 2015 when strains having both N179 and N104 dominate. N179 strains are found in every year. Rarely, in addition to N104 a residue other than N179 or N136 has a potential glycosylation site (grey). Unlike N179, a single mutation with N136 alone is sufficient for glycosylation11. **c,** A/California/07/2009 (PDB: 3LZG) hemagglutinin crystal structure with N179 highlighted in green and the compensatory mutation, I233T, in purple adjacent across the trimer. A slightly earlier antigenic site mutation S101N is shown in red. Inset shows magnified view of the two mutated residues. **d,** HA Western blots of A/California/07/2009 mutants with permutations of N179, K180Q, and I233T with and without PNGase treatment to remove glycans. The single mutant N179 alone does not shift in size, indicating it is not detectably glycosylated. While a portion of the double mutant N179/K180Q shifts, a complete shift, indicating majority glycosylation, is only seen with the triple mutant N179/K180Q/I233T.

### Epistatic changes are necessary for glycosylation

N179 viruses were present at low frequencies in each season since pH1N1 introduction (Fig. 5b), demonstrating that selection for such viruses was not sufficient to overcome fitness limitations in the human population. Viruses with this mutation did not selectively sweep the human pH1N1 strains until the 2015-2016 season (Extended Data Fig. 6b). All of these strains, unlike previous N179 strains, also had an I233T mutation.

The pH1N1 HA structure provides a ready explanation for the epistatic interactions of the HA residues 179 and 233. Residues 233 and 179 are in close proximity across the trimer interface (Fig. 5c), with the change from bulkier and more hydrophobic Ile to Thr, likely facilitating the addition of the large, polar glycan on the other side of the trimer interface. In H3 strains with an N179 glycan, the residue analogous to 233 in pH1N1 is also a small polar residue (Ser).

Residue 179 is located in an antigenic site, called Sa, that is immunodominant in many contemporary human Ab responses. Prior to the N179 glycan sweep, over the 2013-2014 season, pH1N1 was swept by strains antigenically distinct due to a K180Q mutation^28,29^. Surprisingly, through reverse-genetic manipulation of the California/07/2009 strain, we found despite having the amino acid sequence necessary for glycosylation, neither the N179 single-nor N179/I233T-double mutants added a glycan to HA in virions. Rather all three mutations: N179, I233T, and crucially, K180Q were required for full glycosylation of the 179 site (Fig. 5d).

Based on biochemistry and immunology we can reconstruct the selection of N179 glycan addition. Antibody pressure on the Sa antigenic site first selected for the K180Q escape mutation, which replaced other pH1N1 strains. Several years of immune pressure against this new site led to co-selection of the N179 glycosylation with the I233T mutation, which enabled accommodation of the new glycan in the trimer.

### Phylogeographic reconstruction of glycan sweep

A phylogenetic analysis of all HA sequences from pH1N1 viruses collected globally during 2009-2017 revealed that N179 viruses emerged repeatedly during 2009-2014, but did not transmit onward (**Extended Data File 2**). In January 2015, the earliest N179/I233T strains in our database appear in Costa Rica, Taiwan, Turkey, and Iran. Phylogeographic analysis suggests that the N179/I233T clade first emerged in the Spring of 2014, perhaps in the Middle East (Fig. 6, posterior probability 0.8). The virus likely first disseminated from the Middle East to Asia and North America, and then onward to Europe and other regions. By the Spring of 2016, only 15 months after the first 2-glycan strain appeared in the sequence record, previously circulating 1-glycan strains are no longer found. North America had the largest number of N179 viruses, but the inference that N179 strains may have first arose in the Middle East was found to be robust to sample bias (**Extended Data Fig. 8**).

**Figure 6.**
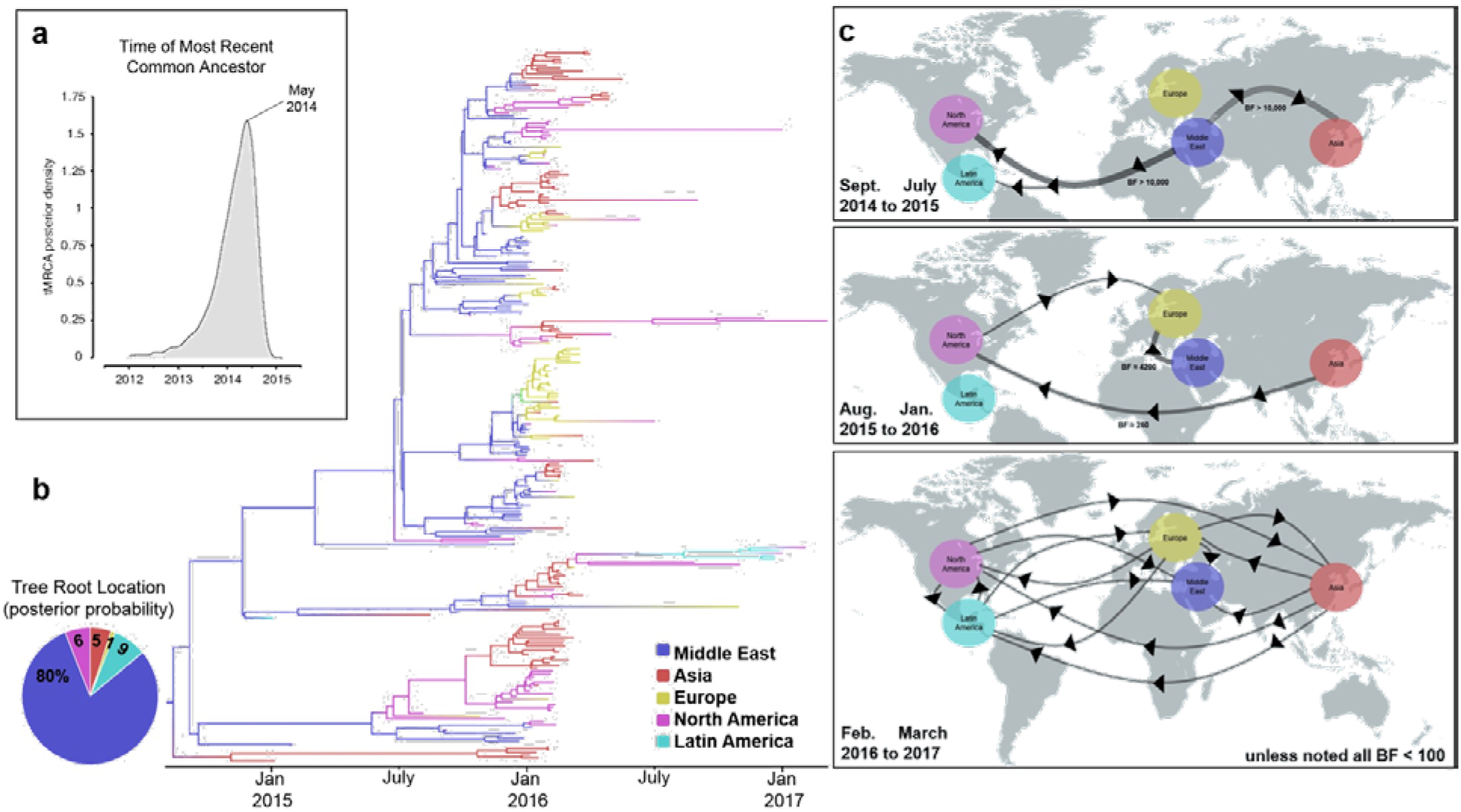
Evolutionary relationships and reconstruction of the dispersal of N179/I233T HA strains. **a,** Posterior density for the time to Most Recent Common Ancestor (tMRCA). Origin is estimated to have occurred in May 2014 (95% Highest Posterior Density: August 2013–November 2014). **b,** Time-scaled MCC tree inferred for 255 N179/I233T HA sequences collected worldwide during 2015 – 2017 (dataset B). The color of each branch indicates the most probable location state. Pie chart indicates posterior probabilities for locations at the root of the phylogeny. The Middle East is estimated as the location where N179/I233T strains emerged with a posterior probability of 0.80. **c,** A possible model for the spatial dispersal of N179/I233T strains was inferred from the MCC tree, based on the data available, and visualized using SpreaD3 at three time points30. N179/I233T strains emerged around May 2014, potentially from the Middle East disseminating to Asia, Latin America and North America, and then globally from February 2016 on. Unless noted on the figure, the Bayes Factors are < 100.

### Forecasts for IAV evolution

Using the three patterns that trace the clock-like evolution of HA glycans, we can forecast future IAV glycan evolution (Fig. 2a and 3a, grey areas). H3 HA reached its glycan limit in 2004, and consistent with Pattern 3 above, swapped a glycan (N144 to N158) during the 2015-2016 season (Extended Data Fig. 6). In addition to this example, when we include all subtypes (H1, H2, and H3), there are 5 additional cases where functionally constrained viruses (i.e. at the glycan limit) adjust glycans or are replaced by a pandemic after 9- to 12-year intervals (Fig. 4). These data suggest H3 will either undergo a second glycan swap or be replaced by a new pandemic HA gene from the animal reservoir over the 2024-2026 seasons.

pH1N1 added a glycan during the 2015-2016 selective sweep. By Pattern 1, another glycan addition should occur during the 2020-2022 seasons. The first 3 glycans added to sH1N1 were N104, N179, and N286 (see strain A/Hickox/JY2/1940). Pandemic H1N1 started with N104, and recently added N179, if pH1N1 recapitulates sH1N1, the next glycan added would be N286.

## Discussion

The rise of N179/I233T over the 2015-2016 season demonstrates how HA glycan evolution can drive rapid selection for strains capable of global human circulation. Through quantitative analysis of trends in HA glycosylation for H1 and H3 viruses over the last century, we found 3 patterns underlie glycan evolution, tracing a predictable life cycle for pandemic HA. Initially, a pandemic HA with few glycans appears. Then, at regular intervals, additional glycans are added, until a functional glycan limit is reached. After reaching this limit, glycans can be swapped, until the highly-glycosylated HA is eventually replaced by a novel, minimally-glycosylated pandemic HA from the animal reservoir.

The consistency of this pattern begs explanation: why are 4- to 6-year intervals required for glycan-addition mutants to become ascendant, and why is twice this interval required to swap between glycan sites after the functional glycan limit has been reached? Part of the answer, no doubt, is the time required for herd immunity to apply sufficient pressure. Another part of the answer appears to lie with the need for epistatic mutations to restore HA function. In any event, our findings can immediately be applied for vaccine selection, since they predict when strains with new glycans will become dominant and identify seasons with higher pandemic potential.

## ACKNOWLEDGEMENTS

This work was supported by the Division of Intramural Research, NIAID. Glennys Reynoso provided outstanding technical support. Jia Jie Wei, Mina Seedhom, and Davide Angeletti provided insightful conversations. Zhiping Ye at FDA generously supplied viruses. SEH is funded through NIAID grants 1R01AI113047 and 1R01AI108686. NST is supported by NIAID’s Centers of Excellence in Influenza Virus Research and Surveillance (HHSN27772201400008C).

## AUTHOR CONTRIBUTIONS

I.K. and M.O.A grew, purified and analyzed viruses. M.A., N.S.T., M.I.N., and M.O.A developed and performed bioinformatic analysis. S.E.H., S.J.Z., and J.S.G. rescued and analyzed glycan mutants. M.I.N, M.A., S.J.Z, and M.O.A. developed the figures. J.Y. and M.O.A. designed and supervised the study, then wrote the manuscript with input from all authors.

## SUPPLEMENTARY INFORMATION

### MATERIALS AND METHODS

#### Virus preparation and purification

We propagated each virus by infecting 5, 10-day old embryonated chicken eggs with 100μL of viral stock diluted 1:2000 in PBS. Infected eggs were incubated at 34.5°C and 65% humidity for ∼41h. We first clarified the allantoic fluid (AF) by centrifuging 4500 x g for 20min, then pelleted the virus by centrifuging for 2h at 26,000 x g. After incubating the viral pellet in 1.7mL PBS overnight at 4°C, we layered the virus onto a discontinuous 15%-60% sucrose gradient and spun at 34,000 x g for 2h. We collected virus at the gradient interface and spun it down a final time in PBS at 34,000 x g for 2h. Virus was resuspended in 100μL PBS and stored as milky opalescent suspensions long term at 4°C. Hemagglutination titers for each virus were measured from neat AF. A complete list of viruses and their origin is found in **File 1**. Total protein content of each virus prep was determined by Coomassie blue-based assay performed per manufacturer instructions [Bio-Rad, DC Assay].

AF from eggs infected with 72 out of the 75 viruses we attempted to grow showed hemagglutination activity (HAU), indicating viral presence. We collected pure virus from all viruses in amounts ranging from 20-6920 μg. Sham infection and purification of AF from control eggs did not produce enough protein for detection, and ran cleanly on a gel (10 μL of a 100 μL prep, Lane #29, Extended Data Fig. 1).

To determine the identity of individual bands on the Coomassie gel, we grew H1N1/H3N2 reassortant viruses A/HK/68 (HK), A/PR8/MCa (PR8), X31 (HK HA and NA, PR8 background), J1 (HK HA, PR8 background), and P50 (PR8 HA, HK background). These viruses were run together on the same gel and immunoblotted with appropriate antibodies to identify which bands corresponded to which proteins via Coomassie blue staining (Extended Data Fig. 2).

To determine the effect of S179N, K180Q, and I233T mutations on glycosylation of pH1N1 HA, we made HA plasmids with different mutation combinations in A/California/07/2009 HA background, which also included an additional D239G mutation to increase yield in eggs (Fig. 5d). Reassortant viruses with mutant HA and NA from A/California/07/2009 and internal segments from PR8 were rescued and grown in eggs. Virus was enriched from AF by centrifugation, standardized by ELISA using a stem reactive antibody (C179), and visualized by SDS-PAGE using the anti-HA CM-1 antibody. A portion of each virus was treated with PNGase F.

#### Protein gels and immunoblotting

We mixed purified virus (1 μg protein) with 4× NuPage loading buffer (Invitrogen), and boiled for 10 min at 96°C. We electrophoresed samples with Chameleon Duo Li-Cor ladder on 4–12% Bis-Tris Gels (Invitrogen) at 200 V for 55 min. To visualize proteins, we fixed gels for 10 min with 10 ml 10% acetic acid and 50% methanol, shaking at RT. After removing fixative, we added 10 ml GelCode stain (Pierce) and shook for 30 min at RT, then destained the gels with water overnight. For immunoblotting, we transferred proteins from gels to PVDF membranes with the iBLOT at P3 setting for 7 min. We blocked membranes for 30 min at room temperature (RT) StartingBlock (Thermo). After incubating with primary Ig or sera for 1 hr at RT, and washing 5x for 5 min each in TBST (10 mM Tris, 150 mM NaCl, 0.1% Tween-20), we added secondary Ig, repeating the washing step incubation.

We imaged blots and Coomassie gels on a Li-Cor Odyssey. Band signals and molecular weights were calculated using Li-Cor’s ImageStudio software. Glycoprotein molecular weights were normalized for gel distortion, commonly called “smiling,” by adjusting each glycoprotein molecular weight by the average amount that the M1 and NP molecular weight for each virus varied from the mean molecular weight of M1 and NP proteins on each gel (Extended Data Fig. 3). Linear regression, Pearson coefficient, and P-values were calculated using PRISM software.

Two data points in Fig. 1 are outliers. The first, A/Georgia/M5081/2012 (#28), a pH1N1 is predicted to have 1 head glycan, but migrates much slower then expected. We made a new viral stock, and deep-sequenced it to confirm its identity. There were no additional predicted N-linked glycans. This HA alone in our panel of H1N1s (which includes another contemporary pH1N1 A/California/07/2009, #27), does not immunoblot with the anti-H1 HA monoclonal CM1 antibody even though the CM-1 peptide epitope used to generate the mAb is present. We suspect the conformation of this protein under SDS-PAGE conditions is somehow altered, changing the HA migration through the gel, and altering its interaction with CM-1. These interesting and perplexing findings will be pursued in future experiments. The second outlier, A/Victoria/361/2011 (#72) is the only H3N2 predicted to have 8 head glycans in our panel. As indicated in the text, H3N2s with more than 7 glycans are not common, indicating possible functional constraints on H3 HA. A possible constraint might be incomplete glycosylation of the virions.

#### Influenza sequences and N-glycosylation prediction

Human HA protein sequences were retrieved from the NIAID Influenza Research Database (IRD) ^31^ through the web site at http://www.fludb.org (accessed April 20, 2017). Sequences with identical strain names were retained only if the majority of members had identical sequences. Sequences were aligned to A/California/04/2009 (H1N1) (FJ966082) with MAFFT v7.305b, and removed if indels existed in relation to the reference or if any ambiguities existed within the HA head. The amino acid numbering of sites important for this work with both H1 and H3 numbering conventions is shown in **Table 1**. Potential glycosylation sites were identified by searching for the NX[S/T]Y motif where X/Y were any amino acid other than proline or by using NetNGlyc 1.0 artificial neural network-based prediction software^24^. The glycan frequencies used for Fig. 2–5 and Extended Data Fig. 5 are found in Supplementary Files 3-8. Stacked glycan plots (Fig. 2a **+** 3a) are similar to a previous publication^32^. For the quantitative analysis of SDS PAGE in Fig. 1 and Extended Data Fig. 4, residues with positive NetNGlyc scores were considered glycosylated “N+”, and residues with only one negative mark were recorded as “N-”. Both conditions were treated as having a glycan present. Glycan predictions with “N-- were counted as negative.

#### pH1N1 phylogenetic analysis

Sequences glycosylated at N179 from 2009-2015 were aligned by ClustalO^33^. An unrooted phylogenetic tree of all 365, N179-glycosylated sequences was made using FigTree^34^. The earliest strains in the same clade as the SWSs were identified as: A/Shiraz/4/2015, A/Shiraz/3/2015, A/Taipei/0021/2015, A/Adana/08/2015, A/Shiraz/6/2015.

To discover any additional mutations co-occurring with N179, we identified all sequences with >99% sequence similarity to the 6 earliest sweeper strains (SWSs) using USEARCH (max ∼17nt differences)^35^. This represented 15% of all FluDB sequences. An unrooted phylogenetic tree of a downsampled set of 370 sequences representing pre-SWS and SWSs (as well as outgroup A/California/07/2009) was aligned in ClustalΩ is shown to the right. From this alignment, the two conserved amino acid changes that define SWSs, S179N and I233T were clearly seen.

To determine time and location of SWSs emergence, all pH1N1 in FluDB from 2009 to present day were collected and annotated with collection country, date, and glycosylation status. We also screened GISAID EpiFlu database for N179/I233T viruses. We down-sampled more than 16000 sequences to ∼200 sequences per year (dataset A), taking into account location diversity and maximum temporal representation per year (total number of sequences = 1853). Dataset A was aligned using MAFFT v7.310^36^. We inferred a phylogenetic tree using the maximum likelihood (ML) method available in the program RAxML v7.2.6^37^ incorporating a general time-reversible (GTR) model of nucleotide substitution with a gamma-distributed (Γ) rate variation among sites. To assess the robustness of each node, a bootstrap resampling process was performed (1000 replicates), again using the ML method available in RAxML v7.2.6.

A smaller dataset B was assembled from the dominant clade with N179/I233T sequences to reconstruct its evolutionary and dispersal dynamics. Dataset B has 255 sequences from 2015 to 2017 with appropriate geographical and temporal representation.

Phylogenetic relationships were inferred using the time-scaled Bayesian approach using MCMC available via the BEAST v1.8.4 package^38^ and the high-performance computational capabilities of the Biowulf Linux cluster at the National Institutes of Health, Bethesda, MD (http://biowulf.nih.gov). A relaxed molecular clock was used, with a constant population size, and a Hasegawa, Kishino and Yano (HKY) 85 model^39^ of nucleotide substitution with gamma-distributed rate variation among sites. For viruses for which only the year of viral collection was available, the lack of tip date precision was accommodated by sampling uniformly across a one-year window from January 1st to December 31st. The MCMC chain was run separately at least three times for each of the data sets and for at least 100 million iterations with sub-sampling every 10,000 iterations, using the BEAGLE library to improve computational performance^40^. All parameters reached convergence, as assessed visually using Tracer v.1.6, with statistical uncertainty reflected in values of the 95% highest posterior density (HPD). At least 10% of the chain was removed as burn-in, and runs for the same lineage and segment were combined using LogCombiner v1.8.4 and downsampled to generate a final posterior distribution of 500 trees that was used in the subsequent spatial analysis^41^.

The phylogeographic analysis considered 5 locations. 4 Asian countries: China, Japan, South Korea and Taiwan; 2 North American countries: Canada and the United States of America (USA); 2 European countries: Czech Republic and Russia; 3 Middle Eastern countries: Iran, Saudi Arabia and Turkey; 3 Latin American countries: Costa Rica and Mexico. A non-reversible discrete model was used to infer the rate of location transitions, along with Bayesian Stochastic Search Variable Selection (BSSVS) to identify those highly significant transitions while improving statistical efficiency^42^. For computational efficiency the phylogeographic analysis was run using an empirical distribution of 500 trees^41^, allowing the MCMC chain to be run for 50 million iterations, sampling every 5,000. Maximum clade credibility (MCC) trees were summarized using TreeAnnotator v1.8.4 and the trees were visualized in FigTree v1.4.3. The worldwide dispersal of N179/I233T glycosylated viruses was visualized using SpreaD3^30^.

Additionally, we performed a sensitivity analyses by down-sampling all oversampled locations in dataset B to 45 sequences per location (dataset C), in order to address if the phylogegraphic estimates described above, were robust to any sampling bias.

**Extended Data Figure 1.**
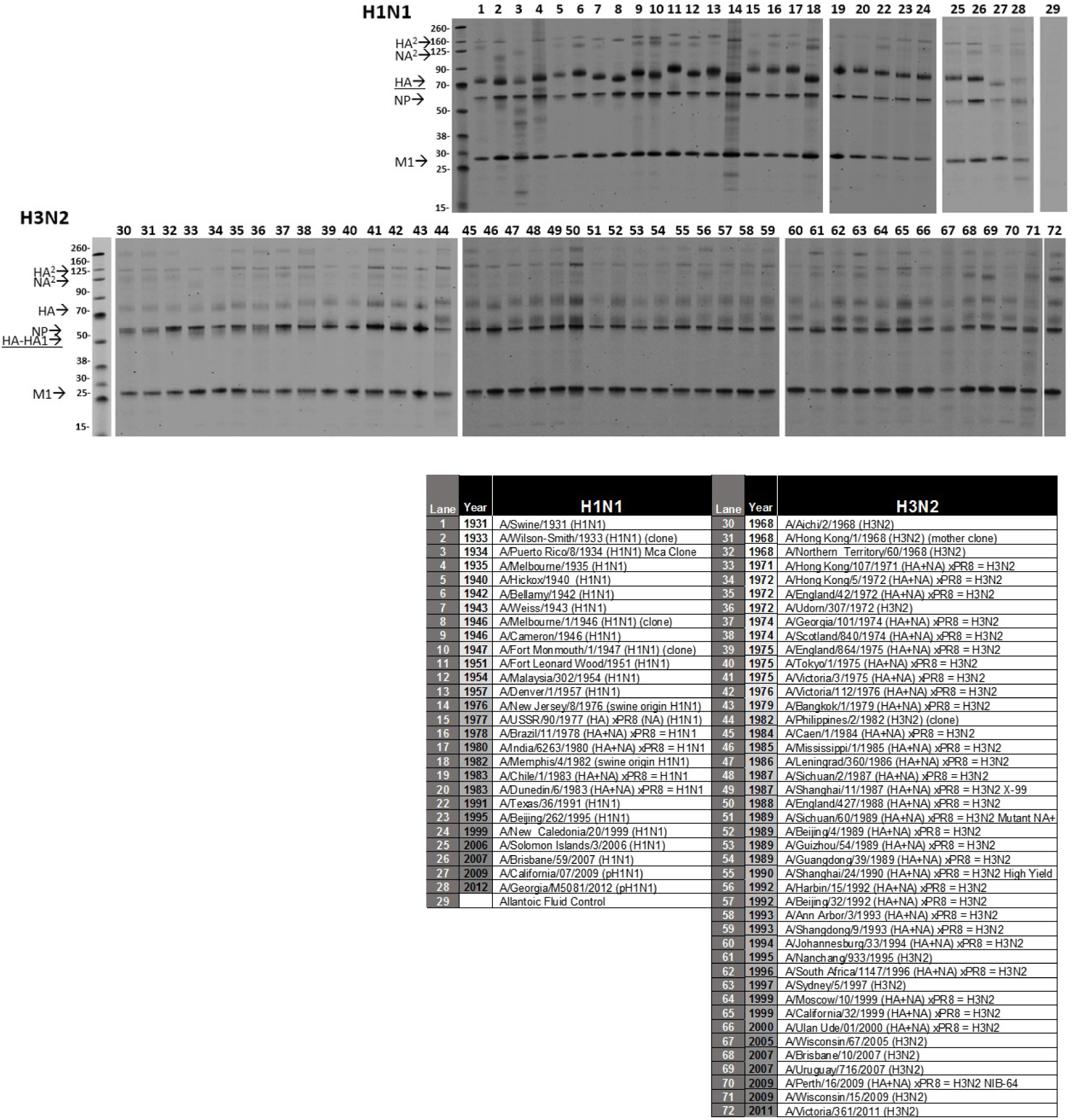
SDS-PAGE of purified virus. Molecular weight standards are labelled. Representative blots are shown. Lane identifications with reassortant status are shown below blots. Source of strain, yield and hemagglutinin units are shown for each lane in **File 1**.

**Extended Data Figure 2.**
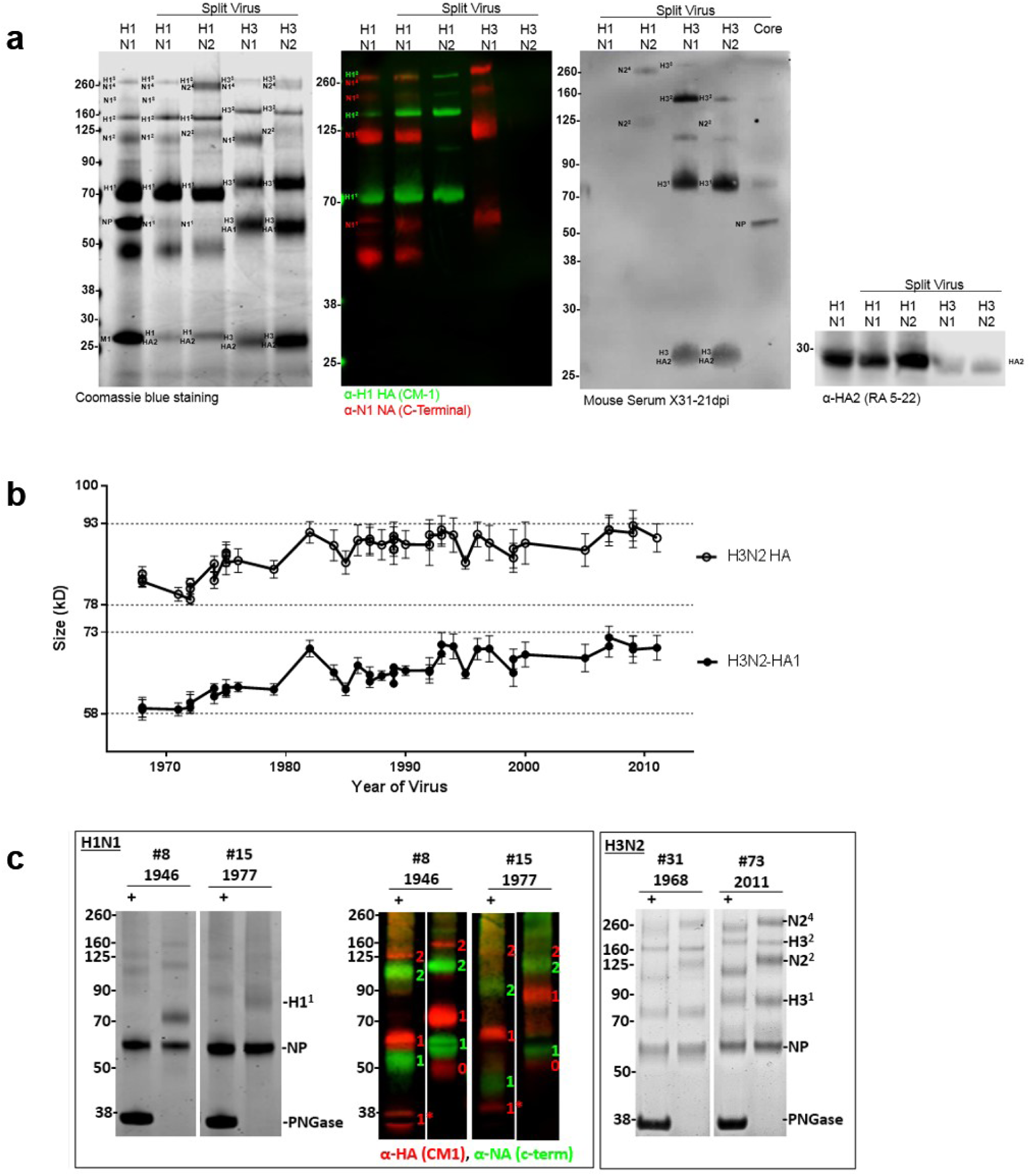
Identification of viral proteins in Coomassie blue stained gels. **a,** Purified whole PR8 or reassortant, detergent split viruses were stained with Coomassie blue; probed with monoclonal antibodies against HA1 (CM1), NA N1 (C-terminal), and HA2 (RA-5 22); or convalescent sera from a H3N2 reassortant infected mouse. **b,** H3N2 HA monomer and H3N2-HA1 band sizes correlate almost perfectly. As the resolution was better for H3N2-HA1, these were used for analysis. **c,** PNGase treatment of viruses shows deglycosylation of glycoproteins for representative viruses. Coomassie stained gels and monoclonal antibodies against H1 (CM1) and N1 (C-terminal) for H1N1.

**Extended Data Figure 3.**
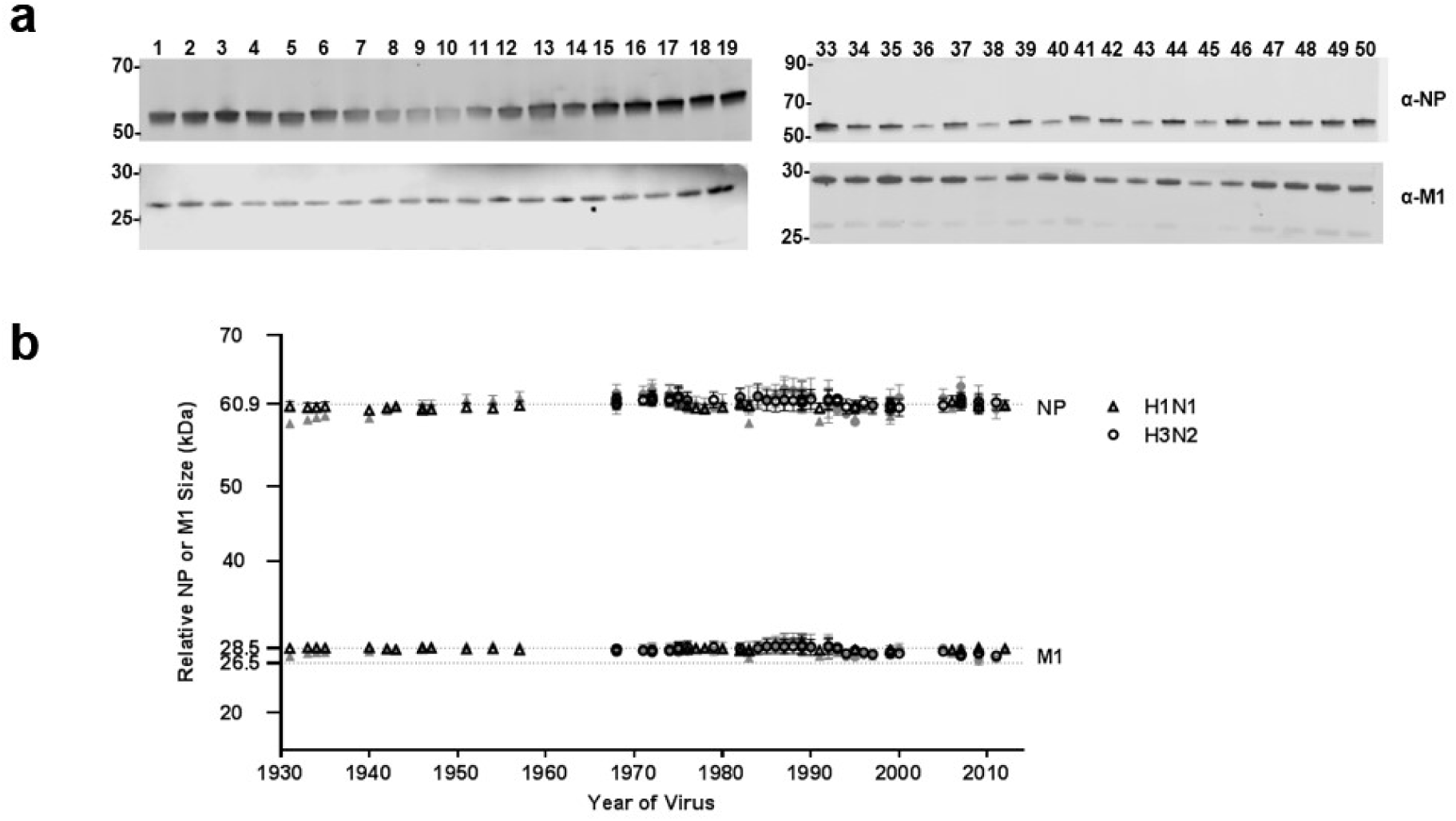
NP and M1 proteins serve as internal ruler. **a,** Western blots of NP and M1 for representative viruses. **b,** Quantitation of NP and M1 relative size from SDS-PAGE. Raw data (grey) was normalized to account for gel distortion (open black symbols). Average size of each protein is shown on the Y-axis. Error bars are standard error from n = 3-6 blots.

**Extended Data Figure 4.**
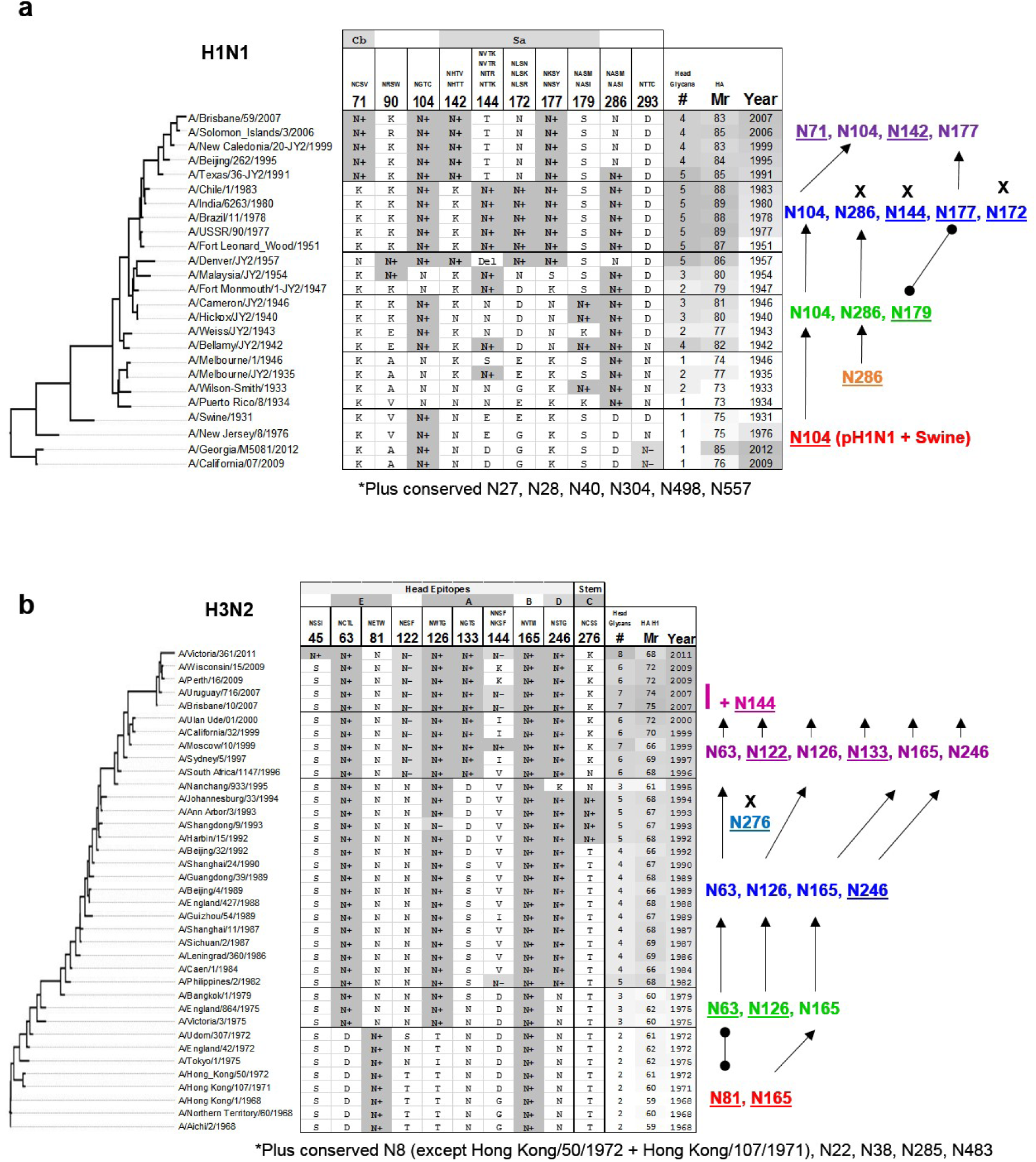
Glycan groups of IAV viral panel. **a,** H1N1 or **b,** H3N2 viruses with the predicted glycosylation status of HA Asn (N+ or N-), the substituted residue present, or a deletion (Del). The total number of head glycans, relative HA size as migration rate through gel (Mr), and year of virus collection is shown for each virus. The glycosylation progression is shown color coded to match individual viruses in Fig. 1b. New glycosylation sites are underlined. Arrows connect conserved glycosylation sites and barbells connect glycans in the same epitope, but not same site. Lost glycosylation sites are noted with an X. Conserved glycans among all viruses are listed below the table.

**Extended Data Figure 5.**
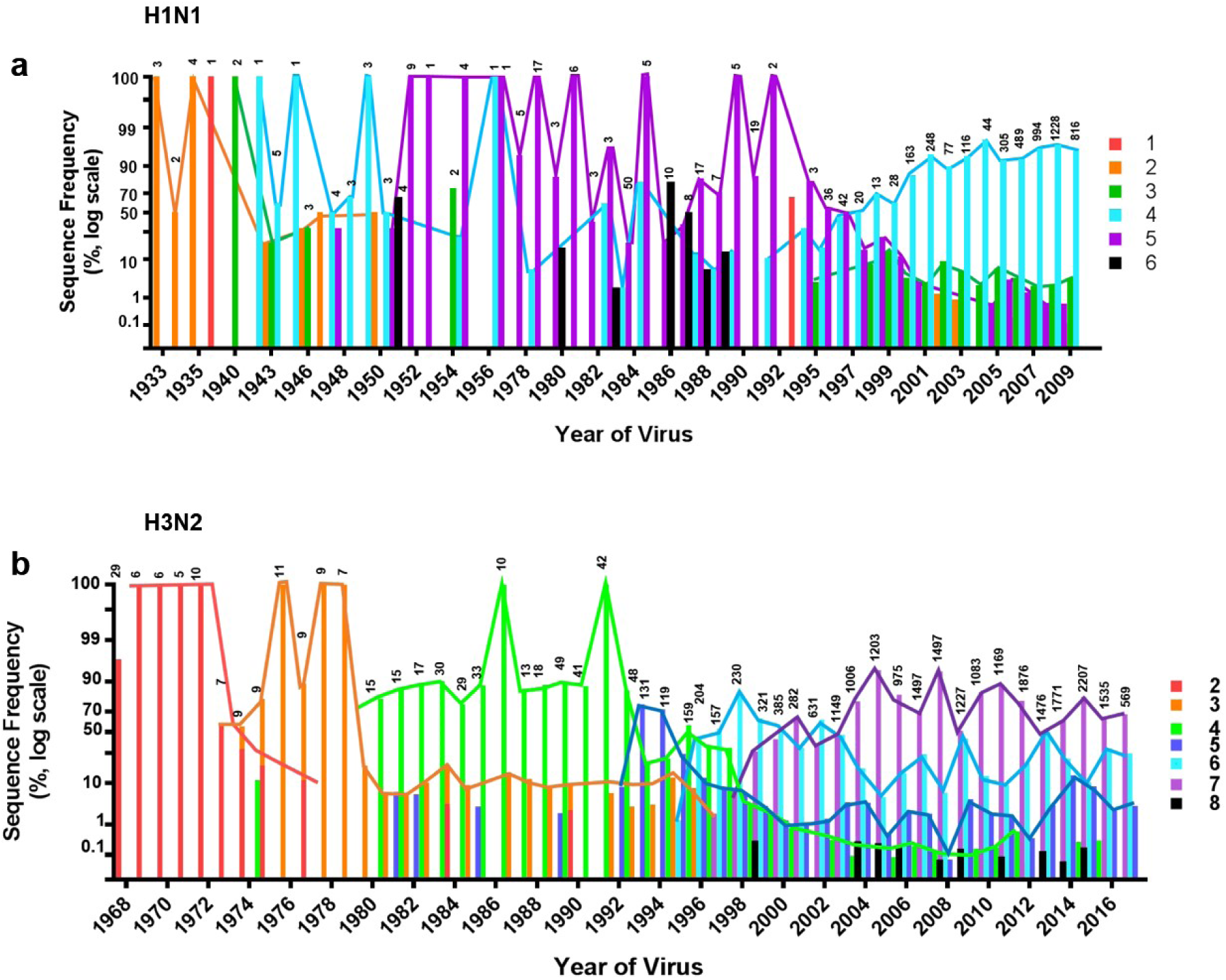
Frequency of total sequences by number of glycosylation sites. **a,** Percentage of H1N1 or **b,** H3N2 sequences in FluDB with indicated number of predicted head glycans. Above each lane are total number of sequences in FluDB from that year. Y-Axis is log scale. Sequences above the functional glycan limit 6 for H1N1 and 8 for H3N2 are shown in black.

**Extended Data Figure 6.**
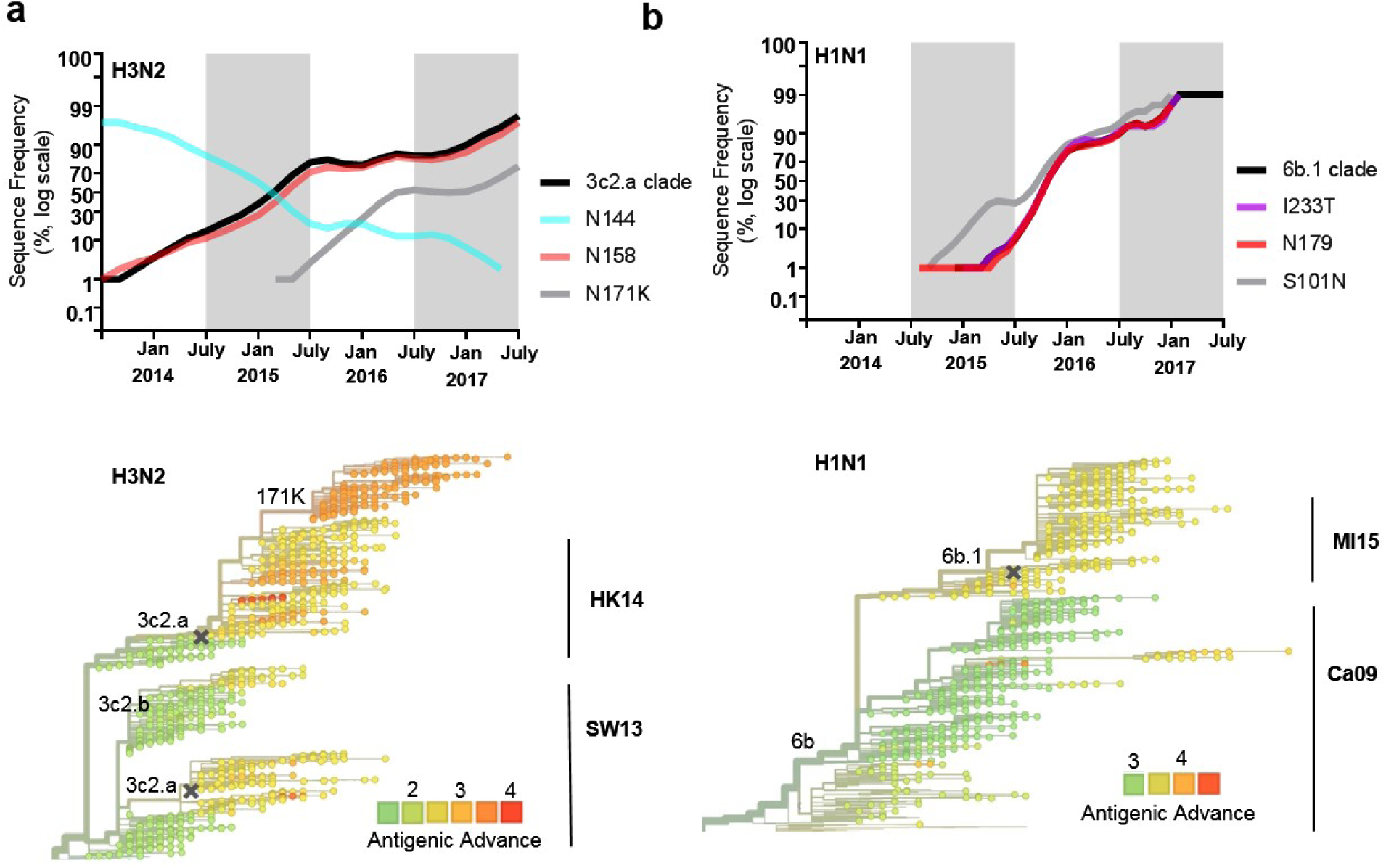
Major IAV clades track with glycan evolution. Using nextflu.org we tracked the most recent glycan changes for **a,** H1N1 and **b,** H3N2 viruses in terms of frequency of the major clade (black) and other mutations of interest. On bottom, phylogenic trees are colored according to antigenic advance measured as log_2_ HI titer distance from the root of the tree. Vaccine strains are marked with an X. Antigenic clusters are labelled to the right.

**Extended Data Figure 7.**
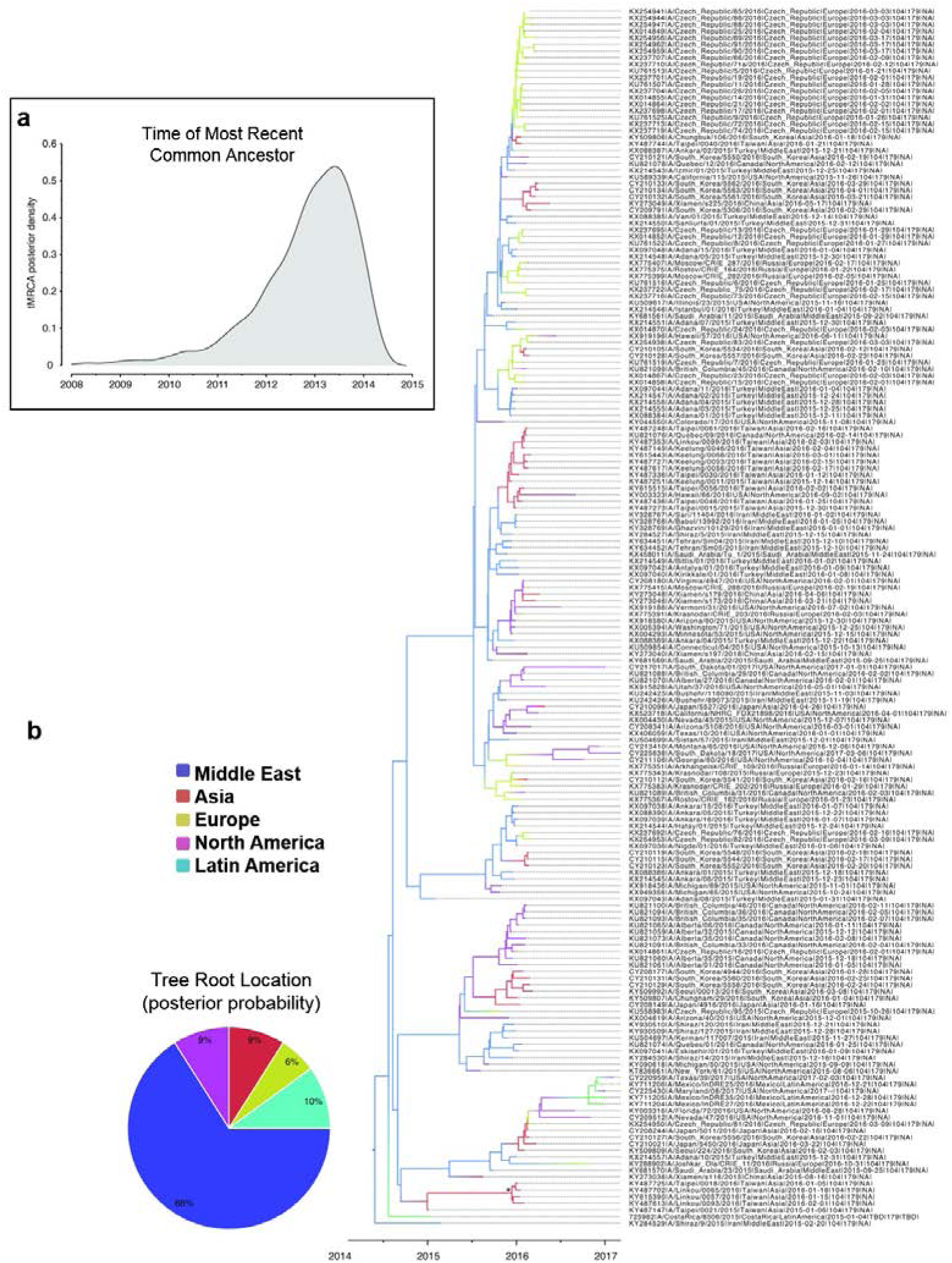
Sensitivity analyses for the evolutionary relationships between N179+ HA sequences. **a,** Posterior density for the time to Most Recent Common Ancestor (tMRCA). Origin is estimated to have occurred in March, 2013 (95% Highest Posterior Density: May, 2011-September, 2014). **b,** Time-scaled Bayesian MCC tree inferred for 184 N179/I233T HA sequences collected worldwide during 2015-2017 (dataset C). The color of each branch indicates the most probable location state. Pie Graph depicts the posterior probabilities for locations at the root of the phylogeny. The Middle East is estimated as the location where N179+ viruses emerged with a posterior probability of 0.66.

**Supplementary Table 1.**
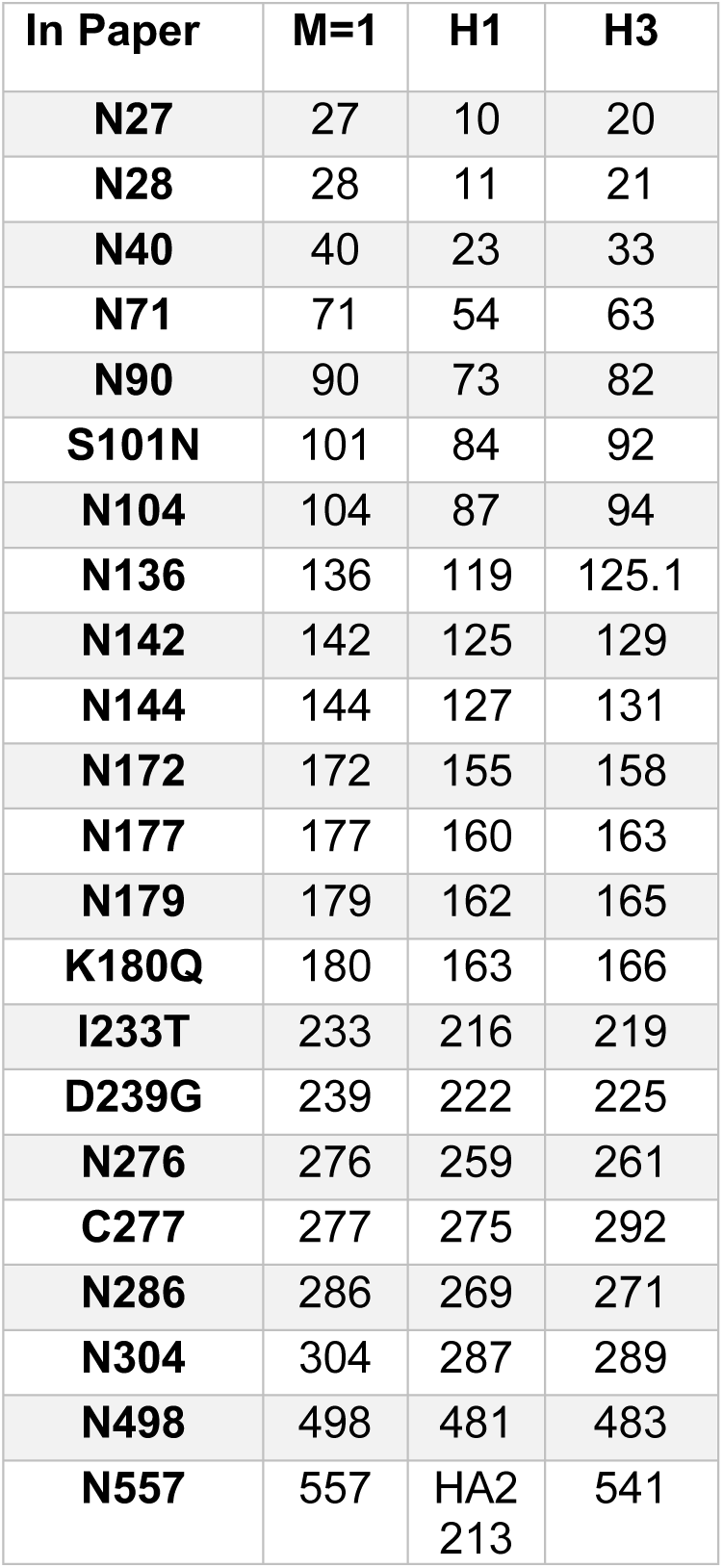
Relevant amino acid residues with H1 and H3 numbering. Calculated using the “HA subtype numbering conversion tool (beta)” on FluDB. Numbering in our text starts from the initial HA methionine (M=1).

| In Paper | M=1 | H1 | H3 |
| --- | --- | --- | --- |
| **N27** | 27 | 10 | 20 |
| **N28** | 28 | 11 | 21 |
| **N40** | 40 | 23 | 33 |
| **N71** | 71 | 54 | 63 |
| **N90** | 90 | 73 | 82 |
| **S101N** | 101 | 84 | 92 |
| **N104** | 104 | 87 | 94 |
| **N136** | 136 | 119 | 125.1 |
| **N142** | 142 | 125 | 129 |
| **N144** | 144 | 127 | 131 |
| **N172** | 172 | 155 | 158 |
| **N177** | 177 | 160 | 163 |
| **N179** | 179 | 162 | 165 |
| **K180Q** | 180 | 163 | 166 |
| **I233T** | 233 | 216 | 219 |
| **D239G** | 239 | 222 | 225 |
| **N276** | 276 | 259 | 261 |
| **C277** | 277 | 275 | 292 |
| **N286** | 286 | 269 | 271 |
| **N304** | 304 | 287 | 289 |
| **N498** | 498 | 481 | 483 |
| **N557** | 557 | HA2<br>213 | 541 |

#### Supplementary Files

**File 1. Viruses used for SDS PAGE.** Viral source, yield and hemagglutinin units for each strain for in Extended Data Fig. 1.

**File 2. Maximum likelihood tree of dataset A**. The phylogenetic tree was inferred using the maximum likelihood method for the dataset A sequences, entailing 1853 H1 sequences from around the globe. The tree is midpoint rooted, all branch lengths are drawn to scale, and bootstrap values >70 are provided for key nodes. N179+ strains are shaded blue.

**File 3.** Frequency of pH1N1 sequences by glycan count.

**File 4.** Frequency of pH1N1 sequences by glycan residue.

**File 5.** Frequency of sH1N1 sequences by glycan count.

**File 6.** Frequency of sH1N1 sequences by glycan residue.

**File 7.** Frequency of H3N2 sequences by glycan count.

**File 8.** Frequency of H3N2 sequences by glycan residue.

